# Moderate Diversity of activation thresholds promotes cooperation in the Threshold Public Good Games

**DOI:** 10.1101/2023.10.30.564720

**Authors:** Gergely Boza, Szabolcs Számadó

**Affiliations:** Institute of evolution, HUN-RES Centre for Ecological Research, Konkoly-Thege Miklós str. 29-33., Budapest, Hungary; Department of Sociology and Communication, Budapest University of Technology and Economics, Egry J. u. 1. Budapest, Hungary; CSS-RECENS, HUN-RES Centre for Social Sciences, Tóth Kálmán u. 4, Budapest, Hungary; Cooperation and Transformative Governance Group, Advancing Systems Analysis Program, International Institute for Applied Systems Analysis (IIASA), Schlossplatz 1, Laxenburg, Austria

## Abstract

Societies face various collective actions which posits social dilemmas, in which a certain number of group members must act cooperatively in order to reach a collective goal. Such social dilemmas are often modelled as Threshold Public Good Games, in which the collective goal is reached successfully if the number of cooperative decisions reaches a threshold. Because cooperation is often a costly act, to coordinate such actions actors can attempt to communicate, to ensure that cooperative decisions are only made once the chances of reaching the goal, thus meeting the threshold is secured. Here we focus on the distribution of activation thresholds in societies, which captures the dynamics of peer pressure and the interaction between different levels of selfish and cooperative behaviours. Here we show that moderate diversity of activation thresholds favours cooperation under wide range of parameters.

## Introduction

All social systems are characterized by sets of dynamic interaction between people and their environment in (Holling 2001; Vespignani 2012). The participation of people in the various aspects of social tasks is shaped by the interplay between the decisions, actions, interactions and reactions of the collective (Holling 2001; Brennen et al. 2016). Hence, the outcomes of collective actions in such behavioural ecosystems (behavioural ecological systems, BEM) are the result of a coevolution (co-creation) in which individual decisions are continuously influenced by the decisions of others (Asch 1955; Hovell et al. 2002; Bond 2005; Vasconcelos et al. 2013; Trenchard-Mabere 2016). It is, therefore, important to understand the motivations and behaviours of people within their social system to construct a model following a behavioural ecological approach (Brennan 2016).

Cooperation has been a central mechanism shaping natural societies, from molecules to animals, and has had a major role in the evolution of the modern human societies (Jenkins 1983; Kollock, 1998; Maynard Smith & Szathmáry 1997). Collective behaviors can have complex driving forces but usually are created through dynamic interactions of actors sharing common goals in a given social situation (Wiedermann et al. 2020). But as numerous aspect of human lifestyle rests on some form of collective cooperation, all can be undermined by a social paradox. Although everyone would be better off with cooperation for self-regarding individuals it is always beneficial to defect, as cooperation is costly. Thus, defection is expected to dominate the outcome instead of collective action (Olson, 1965). As a consequence, without additional fostering, no individual will be rational to contribute and voluntary collective action is doomed to failure (Olson 1965). This is well demonstrated in experimental public goods games where the public contribution is steadily decreasing over time.

Therefor the presence of successful cooperative endeavors seems to be a puzzle. There is a wide range of proposed solutions ranging from selective incentives (Olson 1965; Oliver 1993), different forms of heterogeneities, such as value, interest, or resource heterogeneity, that usually translates into heterogeneity of individual motivations (Marwell et al. 1988; Marwell and Oliver 1993; Centola 2013a, b), or different structural features of the social networks (Granovetter (1973); Macy 1990;Santos et al. 2008; Centola 2013b). While these mechanisms can stop the decline or even increase the public contribution another proximate problem remains. It is not enough for individuals to have the incentive to cooperate (ultimate explanation) they need to coordinate their actions to achieve successful cooperation.

There is a not so hidden coordination problem in all threshold public goods game situations (see Granovetter 1978 for examples). Many forms of collective actions depend on a “critical mass”, a threshold number of participants to provide the majority of contributions for some sort of a good for the whole collective (Granovetter 1978; Oliver & Marwell 1985; Oliver 1993). These have to pay the start-up cost and take the initial risk in order to induce and facilitate the success, the extent, and effectiveness of the collective action (Oliver & Marwell 1985). Critical mass is defined as the minimum amount of contributions or minimum level of cooperation above which the growth of participating becomes self-sustaining (Granovetter 1978, Marwell & Oliver 1993, Centola 2013a).

The activation threshold is a simple, but empirical evidence supported decision rule, and is described as the number or fraction of others adopting an opinion or behavior, in our case cooperation, that triggers the person to join the collective, adopt the spreading/ trending opinion, state, or behavior (Granovetter 1978; Oliver 1984; Latané & L’Herrou 1996; Watts 2002; Watts & Dodds 2007; López-Pintado & Watts 2008; Centola 2013a; MacCoun 2012; Singh et al. 2013; Karsai et al. 2016; Mønsted et al. 2017). Such threshold, or more generally speaking individual orientations to engage in social actions, can differ greatly from person to person (Wilson 2000; Corning & Myers 2002; Watts 2002; Watts & Dodds 2007; López-Pintado & Watts 2008; MacCoun 2012; Van Zomeren 2015; Karampourniotis et al. 2016; Karsai et al. 2016; Besta et al. 2019). People with low thresholds cooperate even if none or a few have indicated cooperative behavior yet, whom can be called initial contributors, initiators, innovators, or instigators (Granovetter 1978; Watts 2002; Watts & Dodds 2007; Centola 2013a; Singh et al. 2013; Karampourniotis et al. 2016, 2019; Karsai et al. 2016). Each contributor hence increases the level of cooperation increasing the chance of triggering further cooperative decisions. The early cooperators, the critical mass, thus can mobilize the rest initiating a chain reaction of cooperation. Conservative strategies, on the other hand, will have higher preference thresholds, and in the extreme will never become active, sometimes called immune individuals, laggards, resistant or stable nodes in the context of social contagion (Granovetter 1978; Valente 1996; Watts 2002; Singh et al. 2013; Karsai et al. 2016; Wiedermann et al. 2020). The vulnerable or conditional strategies lie in between these two extremes (Karsai et al. 2016).

It is easy to see that the distribution or heterogeneity in these personal cooperation thresholds can promote collective action, but only if there are sufficient number of people with very low thresholds. Below the lowest individual threshold, defection will dominate, while above a cascade of cooperative decisions can trigger the collective mobilizing even those who join at the late stages due to their high individual thresholds. The classical start-up problem is how to meet this critical mass threshold of participation and how to get actors join before this is reached.

We model individual decision-making as a dynamical process, in which individuals interactions are embedded in a behavioural ecosystem, such that the decisions of agents are influenced by the decisions of peers through the time period of coordination, and before the time period of collective cooperation (Vasconcelos et al. 2013). In particular, we will focus in individual variations in preferences or activation thresholds (Granovetter 1978; López-Pintado & Watts 2008; MacCoun 2012; Centola 2013b; Karampourniotis et al. 2016, 2019), and how their magnitude and distribution affect the outcome of collective actions.

In the next section we describe the model. After that we present the results. We found that mid-range diversity promotes cooperation in most of the cases. Finally we discuss the relevance of the results and potential extensions of the model.

## Model

We study an agent-based model of collective action with a focus on personal activation threshold functions and their distributions in the society. We model the collective action within the framework of evolutionary game theory, in particular as a non-exclusive and non-rival Public Good Game (Kollock 1998) played in a society with no particular network structure (well-mixed system) in which individuals randomly join small groups for a cooperative endeavour. For the sake of simplicity, here we assume that every individual is determined to cooperate in any particular group, but their decision to act cooperatively, or to contribute to the collective action, thus their *activation* is dependent on how many other group members have their willingness to cooperate expressed.

For this, we model *M* individuals whose personal attributes include the **personal activation threshold (PAT)** (*T*_*x*_) and the **steepness of their decision probability function (STED)** (*D*_*x*_) (Figure 1). We employ stochastic updating, during which all corresponding interactions and steps described below are iterated for *M* times. The main steps include the random group-formation, the group interactions (i.e. signaling intent to cooperate), the game of collective action, the group dismantling, and finally a social learning phase. The interaction step is iterated *τ* times and is composed of the phases of signaling intention to cooperate and revising decision based on the decisions of other group members. Since we do not consider dishonest communication and fake signals, and the activation thresholds remain the same throughout a group interaction step, once an agent decided at step *t* to be cooperative during the collective action, it will remain unchanged until *t* = *τ*, however it can differ between the update steps. The activation of an agent is governed by the decision probability function:

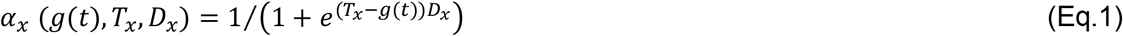

where *g*(*t*) denotes the number of cooperators in the group at time *t* (Figure 1). This formulation allows the investigation of different “archetypes”, such as initiators, laggards, negligent and reluctant. These types differ in their activation thresholds and the steepness of their decision function. The thresholds correspond to deciding early, meaning that only a few players have yet signaled intention, or late during the signaling phase. The steepness corresponds to being determined/rigorous to signal intention and coperate only if the personal threshold is met (steep), or deciding more or less randomly/ making sloppy/casual decisions (flat). We named these types for the shake of discussion, the model allows a continuous variation along these axes.

**Figure 1.**
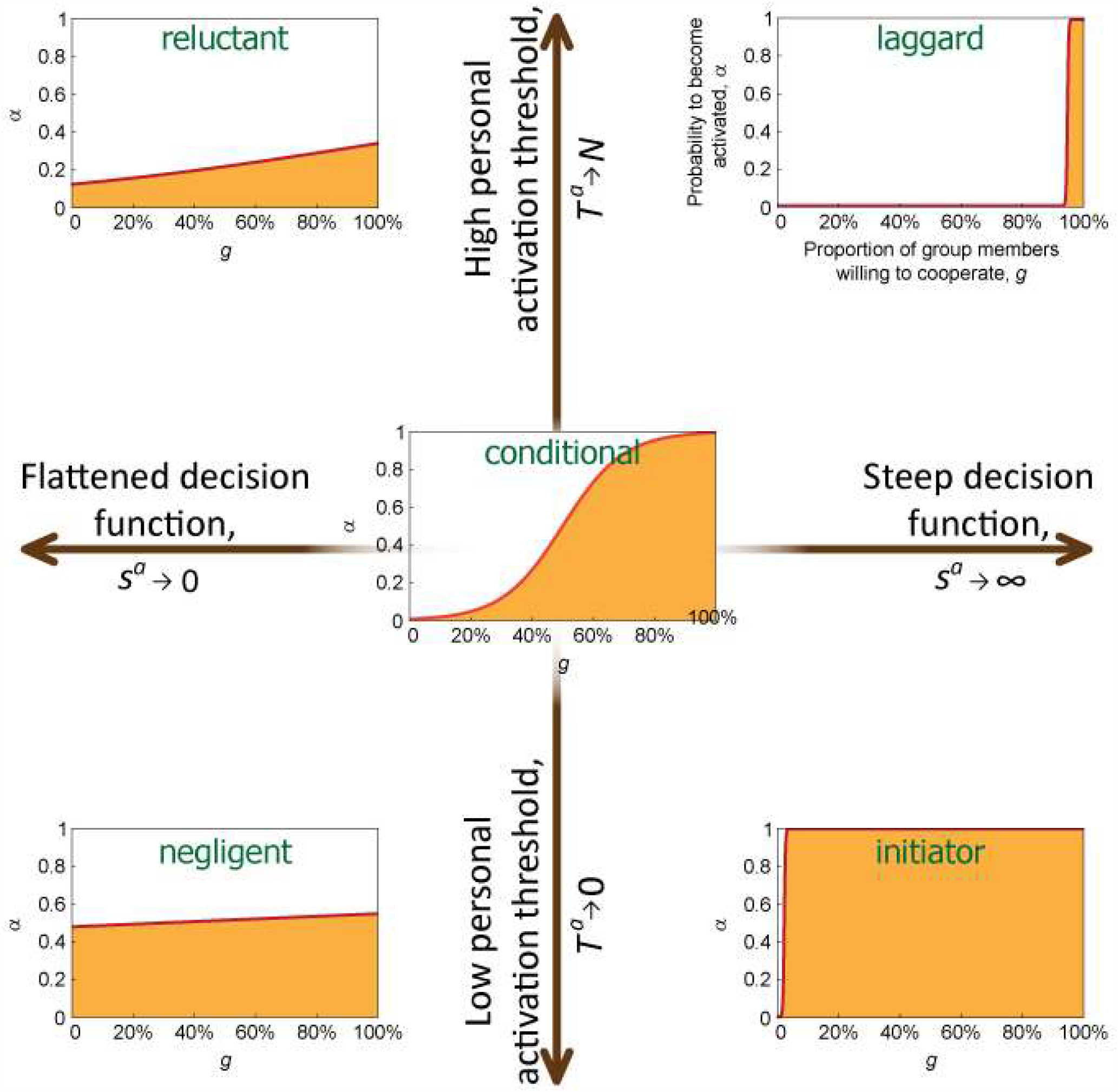
The shape of the activation function depending on the two controlling parameters, the personal activation threshold (*T*^*a*^) and the steepness of the activation function (*s*^*a*^). There is a continuum of different combinations, here we highlighted and named a few strategies.

Naturally, at *t* = 0 *g*(*t*) = 0 for all (newly formed) groups. Once the pre-determined number of interaction rounds are finished (*t* = *τ*), the number of cooperative decisions in the group will determine the success of their collective action.

We model the collective action as a generalized version of the *N*-person Threshold Public Good game or the Volunteer’s Dilemma (Kollock 1998; Archetti & Scheuring 2012), in which the rate of success of the collective action depends on a threshold number (*k*) of cooperators (in general 1 ≤ *k* ≤ *N*, while *k* = 1 for the Volunteer’s Dilemma). The benefit function is defined as:

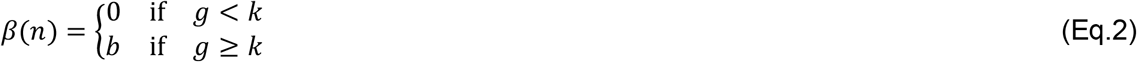

where *b* is the benefit received equally by all group members (Bach et al. 2006; Archetti & Scheuring 2011; Archetti & Scheuring 2012). The payoffs for defectors (D) and for cooperators (C) are written, respectively, as:

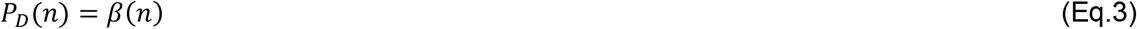

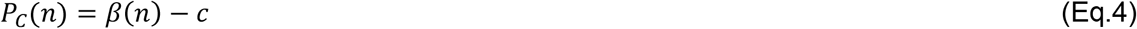

where *c* denotes the cost of cooperation that all cooperators have to pay irrespective of the success of the collective action (Boza & Számadó 2010; Archetti & Scheuring 2012).

Finally, a randomly chosen group member can imitate the more successful strategies through social learning. The focal individual *x* imitates the strategy of a randomly chosen individual *y* from the society, serving as a role model, with a probability *q* according to the Fermi rule, i.e., with probability:

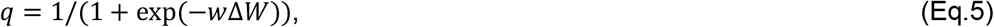

where ∆*W* = *W*_*y*_ − *W*_*x*_ is the payoff difference between the two individuals and *w* specifies the strength of selection.

To model social complexity and behavioral diversity, we assume various distributions of activation thresholds (*T*_*x*_) and types/shapes of decision functions (*D*_*x*_) across the society (Fig. 2.a and Fig. 3.a). The distribution of activation thresholds is modelled as a Gaussian distribution (Granovetter 1978; Watts 2002; López-Pintado & Watts 2008; Karampourniotis et al. 2016; Karsai et al. 2016) cut off at the extreme values, with a mean threshold given as 0 ≤ *μ*^*T*^ ≤ *k*_*max*_ and a variance defined as *VAR*(*μ*^*T*^) = *k*_*max*_*σ*^*T*^ where *k*_*max*_ *= N* + 1. Similarly, the distribution of decision functions steepness is modelled also as a Gaussian distribution, with −*D*_*max*_ ≤ *μ*^*D*^ ≤ *D*_*max*_ and *VAR*(*μ*^*D*^) = *D*_*max*_*σ*^*D*^ where *D*_*max*_ = 100.

**Figure 2.**
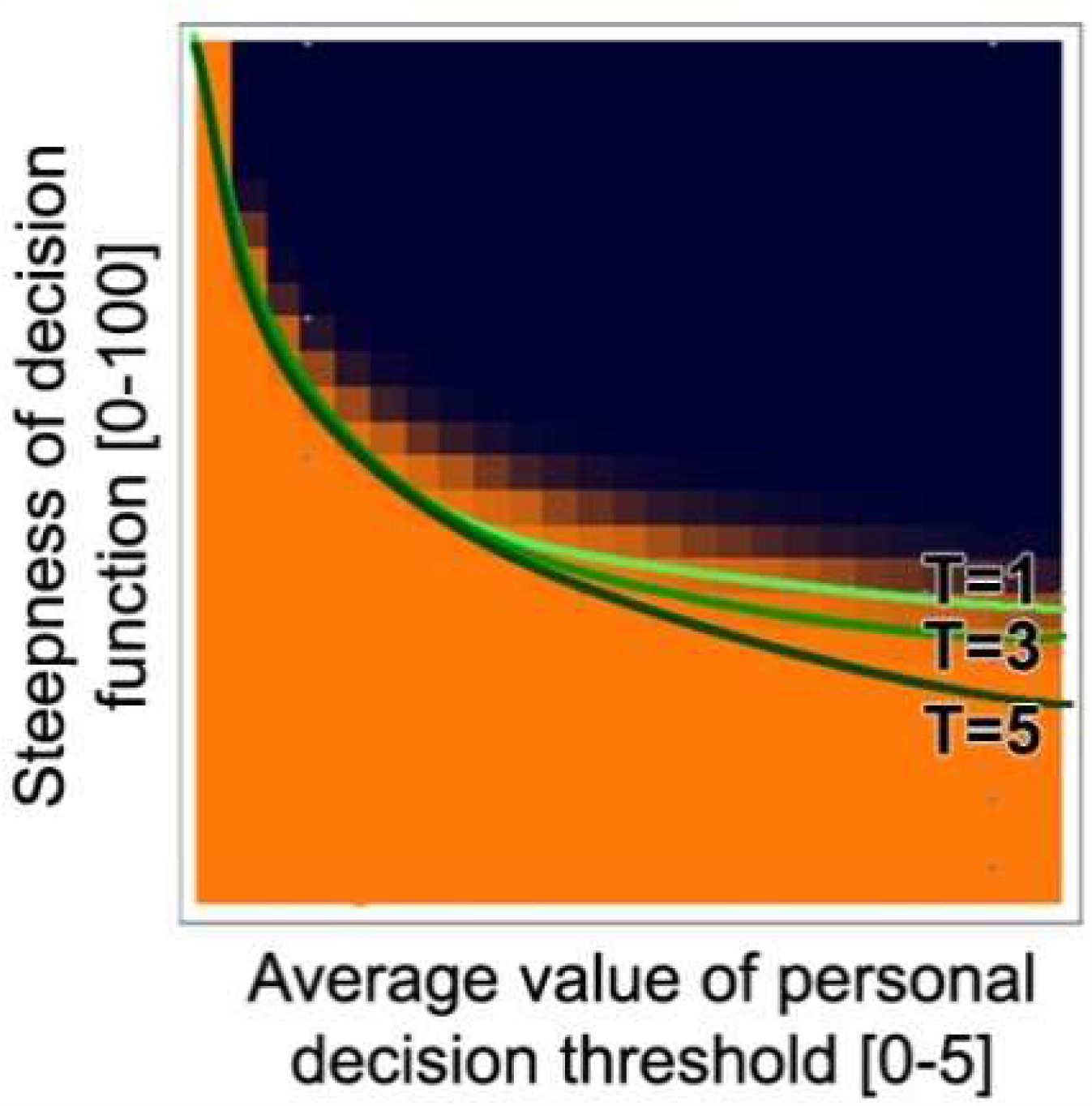
The success of cooperation as a function of the average cooperation threshold (x axis) an the variance of cooperation threshold (y axis). The orange shading depicts a cooperative outcome in collective action, while the black shading depicts failed collective cooperation. Flat decision functions lead to players deciding to cooperate more-or-less randomly, hence the level of personal activation threshold has little effect on the outcome. When the decision function is steep, the collective action is only successful, when the personal activation threshold is low. This is because the bandwagon effect can only start if there are initiator type of players with low activation threshold.

**Figure 3.**
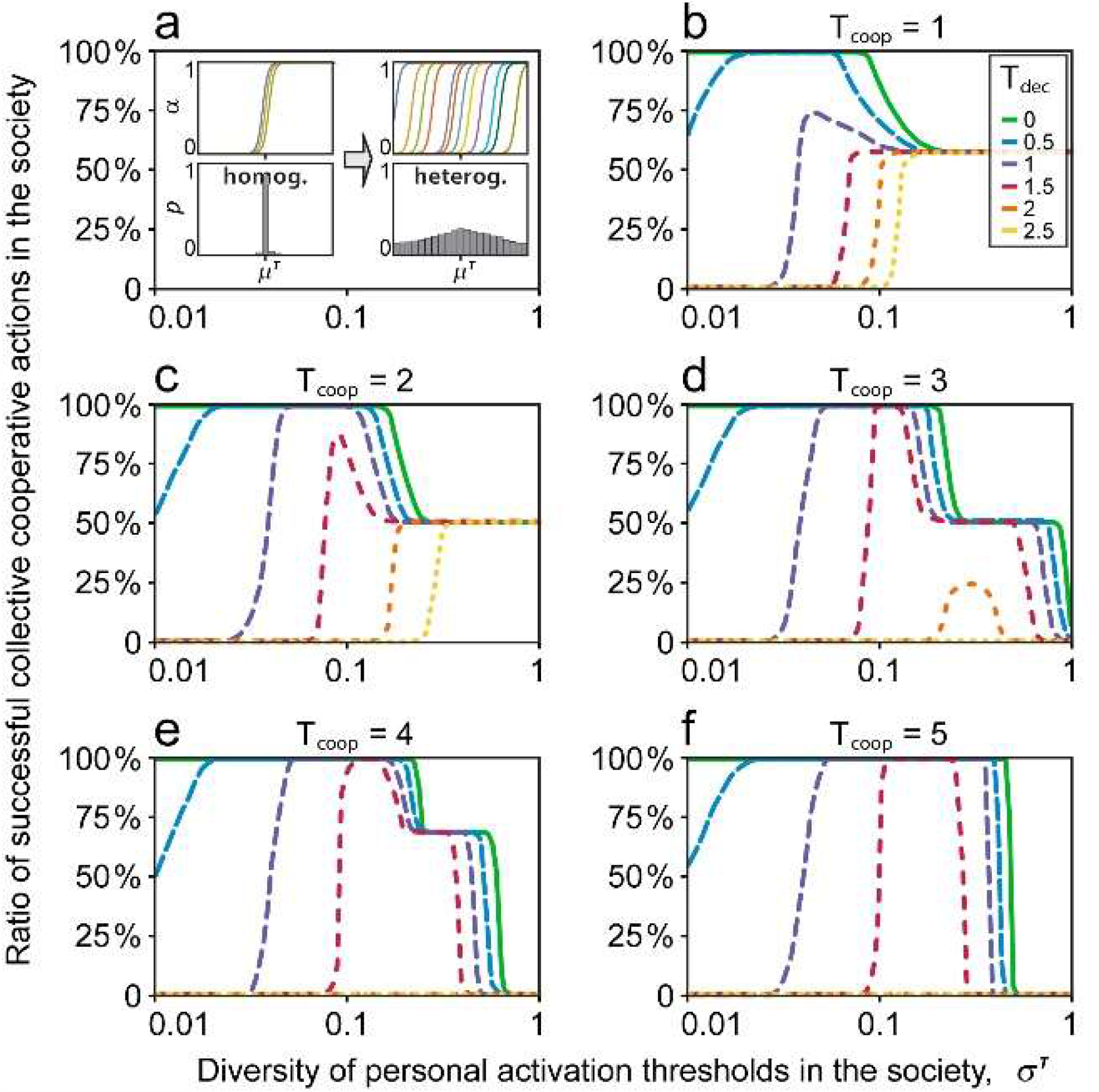
The success of cooperation as a function of the variance of personal decision threshold (x axis).

## Results

The social dynamics in our model system, in general, produces three classes of results: collective failure, universal collective success, and partial collective success in collective cooperation action (see Figures 3,4,5). We investigated the effect of changing the variance of personal activation threshold (PAT) (*T*_*x*_) and the steepness of their decision probability function (STED) (*D*_*x*_) while keeping the other function constant respectively (Figure 3 and 4). Figure 5 shows the interaction of these two functions for every given threshold in a 5 person threshold PGG.

**Figure 4.**
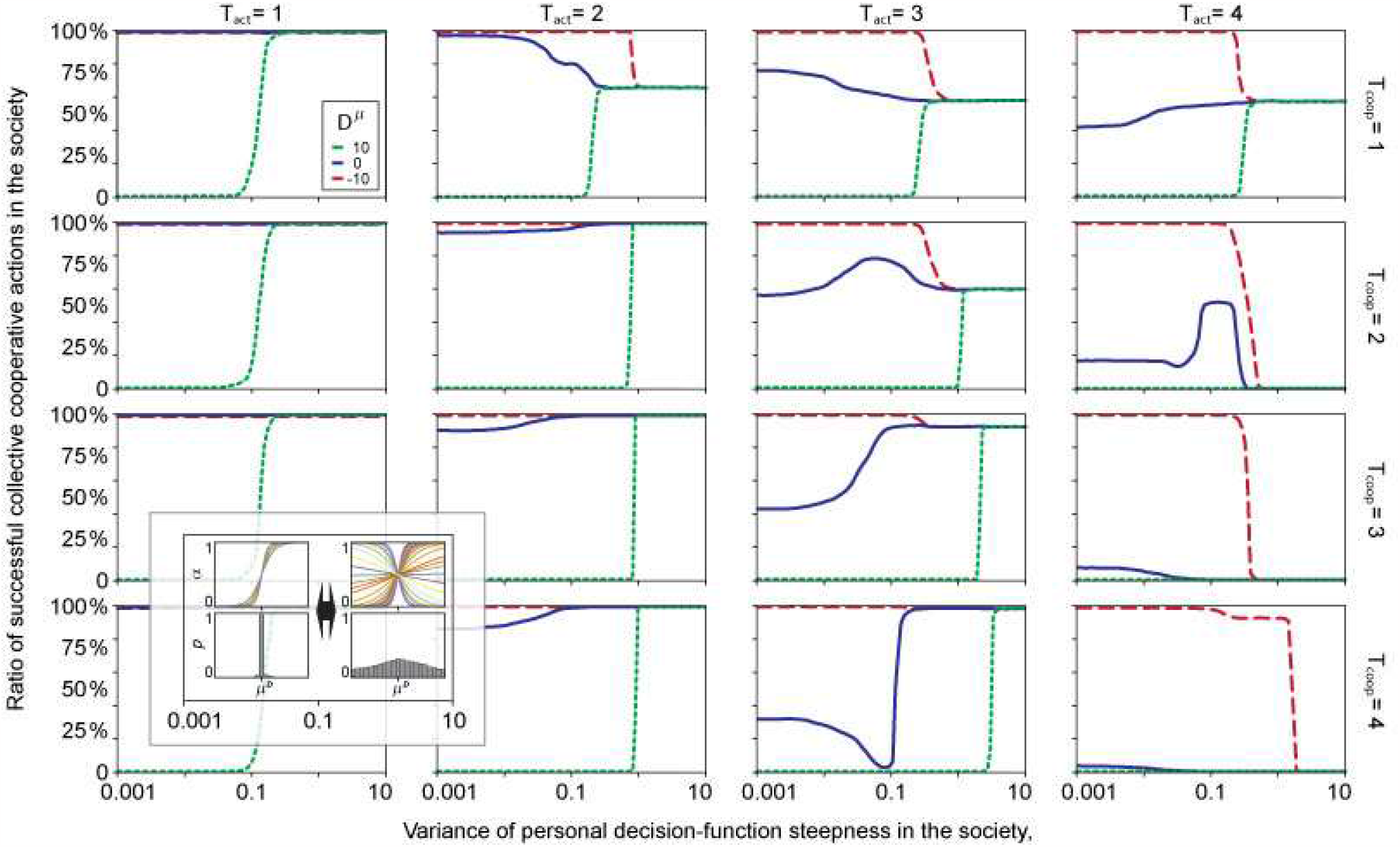
The success of cooperation as a function of the variance of the steepness of personal decision functions (x axis).

The general conclusion is that mid-range variance of PAT favors the highest level of cooperation (Figure 3). Medium to high levels of diversity leads to situations in which success happens only in a certain proportion of the cases. This is because of the random group assembly, groups may form that do not contain players with low activation thresholds (Fig.3 T=1,2,3). Too high levels of heterogeneity (high values of variance) lead to a bi-modal distribution in activation thresholds (since we cut-off at the extreme values), having two peaks at min (0) and max (5) values. In such cases, a few players may join initially, but since the middle range is scarce, the late joiners would never join. Hence, for high values of cooperation thresholds, very high diversity is contra productive (Fig.3 T=4,5).

The interaction of the steepness of their decision probability function (STED) (*D*_*x*_) and variance of STED is more complex (Figure 4). When the decision is conservative (D positive, green dotted lines, Fig.4), meaning that players decide to signal the intention and to cooperate when the personal activation threshold is reached, medium to high levels diversity is needed. The higher the activation threshold, the higher the diversity needs to be to get at least one player to initiate the bandwagon effect. Even this may not be enough, for high cooperation thresholds with high personal activation thresholds, there is not enough diversity to get a successful collective action.

When decision are anti-social (D negative, dashed red lines Fig.4), meaning that players decide to cooperate when the number of other is below their personal activation (or in this case personal de-activation) thresholds. In these cases, low diversity is good. At too high diversity, too many may decide to cooperate, and hence the anti-social players may start deciding to withdraw their signals and intentions and decide not to cooperate after all. Therefore, is such cases, collective action is successful only up to mid-range diversity (high T_act_ values, Fig.4).

Deflated/Flattened decision function (blue lines, Fig.4), which translates into a permissive/ sloppy/random decision function regarding the number of joined cooperators and one’s personal decision threshold, promotes successful collective cooperation for almost all personal decision threshold values (except for high T values, Fig.4). The reason is simple, decisions are more random than are determined by the values of cooperators. Once a few random cooperative decision are made, the bandwagon effect may be triggered, and other will continue to join.

At very high variance of STED the results are the same (Fig.4), because of the distributions are the same regardless of the starting slope.

## Discussion

We employed a model, in which the so called ‘activation threshold’ the level of conditionality of cooperating in a collective actions varied across numerical experiments. In our model the thresholds of collective action (cooperation) and the thresholds of personal activations interact, giving rise to a much more complex dynamics. We showed that a diversity of thresholds is critically important when the number of initiators is not enough to trigger a cascade, in agreement with previous studies of threshold driven contagion dynamics (Karampourniotis et al. 2016). Moderate diversity of PAT and STED is the most efficient promoting cooperation under most of the parameter range investigated.

High diversity is not beneficial most of the time. Previously, Macy (1990) showed that an initially homogenuous population can diverge into different types, “volunteers” and “free-riders”, depending on the production function of the public good. Similar divergence can be observed in our model when the variance of PAT is high. The group with very low PAT joins cooperation early on (“volunteers”), while the group with high PAT never joins (“free-riders”).

Previous works based on the activation or adoption threshold model focused on opinion dynamics, information cascades, culture and language evolution (Granovetter 1978; Castellano et al. 2009; Singh et al. 2013; Wiedermann et al. 2020). Here we used a continuous threshold concept as a way of coordination in the Threshold Public Good Game.

Another difference is that our model considers a probabilistic activation function, in contrast to most of the previous works in which the personal activation threshold was modelled as a (strict and deterministic) step-wise function (Wiedermann et al. 2020).

It has been argued that the egocentric social environment (or social network) plays the most crucial role in an agents’ decision making (Asch 1955; Macy1991; Latané & L’Herrou 1996; Bond 2005; West et al. 2006; MacCoun 2012; Centola & Baronchelli 2015, Ureña et al. 2019.). Groups are formed randomly in our model, this can be appropriate assumption for a large well-connected population. Introducing a social network, thus limiting our agents to team up with anyone, can be an interesting next step.

Peer pressure dynamics is another interesting prospect. Agents who joined the action could pressure the laggards to join. Some argues that such pressure and agents’ resistance to it, is crucial to understand the essence of coordination dilemmas (Karampourniotis et al. 2019).

## Acknowledgements

S.S., and G.B was supported by the National Research, Development and Innovation Office – NKFIH (OTKA) grant K 132250, and by the European Research Council (ERC) under the European Union’s Horizon 2020 research and innovation programm (grant agreement No 648693). S.S. and G.B. acknowledges support from by the National Research, Development and Innovation Office under grant number GINOP-2.3.2-15-2016-00057. The funding agencies had no role in the study design, analyses, or publication.

## Author contribution

S.S. and B.G. conceived the idea, B.G. wrote the code, B.G. and S.S. analyzed the results, B.G, and S.S. wrote the paper.

